# Peer presence elicits task-independent changes within and beyond the mentalizing network across children and adults

**DOI:** 10.1101/2022.04.19.488718

**Authors:** Leslie Tricoche, Denis Pélisson, Léa Longo, Eric Koun, Alice Poisson, Jérôme Prado, Martine Meunier

## Abstract

There is ample behavioral evidence that others’ mere presence can affect any behavior in human and non-human animals, generally facilitating the expression of mastered responses while impairing the acquisition of novel ones. Much less is known about i) how the brain orchestrates the modulation of such a wide array of behaviors by others’ presence and ii) when these neural underpinnings mature during development. To address these issues, fMRI data were collected in children and adults alternately observed and unobserved by a familiar peer. Subjects performed two tasks. One, numerosity comparison, depends on number-processing brain areas, the other, phonological comparison, on language-processing areas. Consistently with previous behavioral findings, peer observation facilitated both tasks, and children’s improvement was comparable to adults’. Regarding brain activation, we found virtually no evidence of observation-driven changes within the number- or language-related areas specific to each task. Rather, we observed the same task-independent changes for both numerosity and phonological comparisons. This unique neural signature encompassed a large brain network of domain-general areas involved in social cognition, especially mentalizing, attention, and reward. It was also largely shared by children and adults. The one exception was children’s right temporoparietal junction, which failed to show the observation-driven lesser deactivation seen in adults. These findings indicate that social facilitation of some human education-related skills is i) primarily orchestrated by domain-general brain networks, rather than by task-selective substrates, and ii) relatively mature early in the course of education, thus having a protracted impact on academic achievements that may have heretofore been underestimated.

**Highlight:**

- Basic math and reading skills were measured in children and adults.
- Participants were alternately observed and unobserved by a familiar peer.
- Behavior showed that children were as facilitated by peer observation as adults.
- fMRI data showed task-independent, observation-driven changes in mentalizing, attention, and reward brain regions.
- All adults’ neural changes were also found in children, except one located in right temporo- parietal junction.

## 1. Introduction

The presence of an observer affects behavior. Generally, it facilitates simple or mastered responses while impairing complex or novel ones (Zajonc). A long history of psychology research has demonstrated the ubiquity of this social facilitation-inhibition phenomenon across species, observers, and behaviors (Bond and Titus; Guerin; van Meurs et al.). We share this fundamental form of social influence with many -if not all- other animal species, including other primates such as macaques (Reynaud et al.), but also songbirds (Vignal, Mathevon, et al.) or drosophilas (Chabaud et al.). Actual observers embodied by friends (Ruddock et al.), strangers (Guerin), or humanized robots (Woods et al.) trigger social facilitation-inhibition, but virtual observers (Miyazaki), or even imagined ones (Hazem et al.), can induce it as well. All behaviors can be changed by others’ presence (positively or negatively, depending on the difficulty of the task they are embedded into), including, in particular, the very eye movements and attention mechanisms that guide vision, our primary window to the world (Liu and Yu; Tricoche, Ferrand-Verdejo, et al.; Huguet et al.; Wykowska et al.). In contrast with this wealth of behavioral data, there is limited knowledge on the neural mechanisms orchestrating others’ presence effects on such a wide variety of behaviors in so many species (Belletier et al.; Monfardini, Reynaud, et al.). Even less is known about the emergence of the neural correlates of social facilitation-inhibition in children, although others, especially peers, might be particularly important during development (Somerville; Steinberg).

Social facilitation-inhibition is a lifelong phenomenon detectable as early as one year of age in humans (Pearcey and Castro). Therefore, understanding the neural correlates of social facilitation-inhibition during development has potential relevance to several domains including childhood obesity (Higgs and Thomas), adolescent risk-taking (Telzer, Rogers, et al.), and education (van Duijvenvoorde et al.). Because peers’ influences can have staggering dramatic consequences on adolescents’ health and life, much of the effort has heretofore focused on understanding the neural correlates of peers’ influences on adolescent decision making (Hartley and Somerville). Interest is nevertheless emerging for the exploration of peers’ influences on cognitive skills relevant to education (Dumontheil et al.). Given peers omnipresence at school, a better understanding of peer presence effects on cognition could provide useful insights to educators about when to minimize, or on the contrary, maximize peers’ presence during learning in order to improve academic achievements.

At least two different neural mechanisms may explain social facilitation-inhibition remarkable ubiquity across ages, behaviors and species. One is that others’ presence might modify neural activity in task-specific networks. All animals having congeners, it could indeed be adaptive for evolution to endow every neural system, whether sensory, motor or cognitive, immature or mature, with some capacity to process relevant social information (Ferrari et al.). Several lines of evidence from research in non-human animals support this hypothesis. In monkeys, for example, a congener’s presence changes activity in the fronto-parietal network subserving an attentional task (Monfardini, Redoute, et al.), but also in the dorsolateral prefrontal neurons encoding a visuo-motor task (Demolliens et al.). In songbirds, a congener’s presence affects early gene activation in auditory areas when the bird is listening and in motor areas when the bird is singing (Woolley et al.; Riters et al.; Vignal, Andru, et al.; Menardy et al.; Hessler and Doupe; Woolley). Some human neuroimaging data are also compatible with the idea that peer presence affects task-specific regions. Being observed changes activity in the (adult’s) inferior parietal region controlling object grasping during a fine grip motor task (Yoshie et al.), whereas it affects the (adult’s and adolescent’s) dorsolateral prefrontal region controlling relational integration during a complex reasoning task (Dumontheil et al.). In adolescents, being observed by a peer enhances the pleasure of risk-taking and the associated activity in the ventral striatal region controlling reward processing (Albert et al.; Chein et al.; Van Hoorn et al.). Thus, several studies in human and non- human animals support the hypothesis that peer-presence effects on behavior might be mediated by task-specific neural systems.

Another possibility, however, is that others’ presence exerts its influence via one or several domain- general neural systems irrespective of the task. This could especially hold for primates, whose brain is thought to include several domain-general networks dedicated to processing social information (Rogier B Mars et al.). Specifically, social presence effects in humans have been associated with our species’ outstanding mentalizing abilities (Hamilton and Lind). Mentalizing is the ability to infer others’ states-of-mind, such as their desires, intentions, or beliefs (Frith and Frith; Blakemore). Explicit mentalizing is generally considered to mature at around 4 years of age, but implicit mentalizing is present by 15 months of age (Kovács et al.). The core mentalizing network identified in the brain across a variety of tasks and stimuli includes four regions: the medial prefrontal cortex (mPFC), the temporo-parietal junction (TPJ), the precuneus/posterior cingulate cortex (PreC/PCC) and the middle temporal gyrus (MTG) (Preckel et al.; van Veluw and Chance). This network is developing early in life, before school age, and is relatively stable across late childhood, adolescence, and adulthood (Fehlbaum et al.; Richardson et al.). Changes in one or more nodes of this network have been reported in several neuroimaging studies that investigated the effects of peer-presence on various behaviors. This holds true for adolescents observed while taking risks (van Hoorn et al.; Chein et al.; Telzer, Ichien, et al.), making prosocial decisions (Van Hoorn et al.), or engaging in complex reasoning (Dumontheil et al.), as well as for adults observed during risk-taking (Beyer et al.), skilled motor performance (Chib et al.) or embarrassing failures (Müller-Pinzler et al.). It is therefore possible that humans mentalize about the thoughts of the observer, even when such mentalizing is not explicitly required. This would be associated with the systematic recruitment of the brain mentalizing network, irrespective of the task at hand. It is interesting to note that some animal data are also compatible with the idea that neural changes produced by a congener’s presence extend beyond task-specific substrates. Chicks, for example, show a dissociation of the brain regions controlling foraging *vs.* the social facilitation of foraging (Xin et al.). Also, despite its 100,000-neuron brain, the drosophila’s uses two distinct brain networks to encode long-lasting olfactory memories, one when flies are tested alone, the other when they are tested in the presence of other flies (Muria et al.). Overall, studies in humans and non-human animals may also support the hypothesis that peer-presence effects on behavior are mediated by a domain-general neural system such as the mentalizing network.

Critically, the two possible neural accounts of social facilitation-inhibition make different predictions when different tasks have non-overlapping neural substrates. According to the task-specific theory, the effect of peer presence on brain activity depends on the task at hand and is localized in task-specific brain regions. According to the domain-general theory, the effect of peer presence on brain activity is independent of the task at hand and is localized in a domain-general network (such as the mentalizing network). To the best of our knowledge, such a paradigm with two different tasks was used in only one previous neuroimaging study of peer presence effects (Smith et al.). Adolescents (15 to 17-year-old) alternatively performed a gambling, risk-taking task, and a go/no-go, response inhibition task, either unobserved or under the belief that an anonymous peer was watching. The go/no-go task did not activate, however, the typical brain substrates of response inhibition, making a cross-task comparison difficult. So, a successful two-task comparison of peer presence effects neural underpinnings is still lacking.

To test between the task-specific and domain-general accounts of social facilitation-inhibition, we used here functional magnetic resonance imaging (fMRI) to measure the effect of peer presence on the neural mechanisms underlying two basic tasks in children and adults: numerosity comparison and phonological comparison. Both tasks are foundational to humans’ math and reading skills (Phillips et al.; Starr et al.). Numerosity comparison consists in comparing quantities using approximate representations of numbers without relying on counting or numerical symbols (Dehaene). It is an early-developing numerical skill, detectable as early as 6 months of age, that has been found to predict mathematics achievement (Starr et al.; Hyde et al.). Phonological comparison consists in comparing the sound structure of words (Phillips et al.). It is an early-acquired language skill, taught in preschool (Qi and O’Connor), that predicts reading achievement (Ehri et al.). In both the developing and the mature brain, numerosity comparisons involve brain areas supporting the representation of magnitudes in the intraparietal sulcus and posterior superior parietal lobule, while phonological comparison involves language-related areas in the inferior frontal and the middle temporal gyri (Prado, Mutreja, Zhang, et al.; Prado, Mutreja, and Booth). In addition, these two types of comparison are at least as much facilitated by the presence of a peer in 8 to 10-year-old fourth-graders as in adults (Tricoche, Monfardini, et al.).

The fact that both numerosity and phonological comparisons are sensitive to peer presence, despite distinct neural substrates, makes them well suited to test between the task-specific and domain-general accounts of social facilitation-inhibition. Furthermore, testing children and adults provides us with the opportunity to evaluate whether the neural mechanism underlying effects of peer presence change with age and expertise with the task. Here we compare 10 to 13-year-olds to young adults to determine whether the neural correlates of peer presence effects are already mature during the pivotal period between elementary and middle school, i.e., between the end of childhood and the beginning of adolescence. Neural changes elicited by familiar peer presence were analyzed across tasks and ages using regions-of-interest (ROI) analyses to assess changes within task-specific substrates, as well as whole-brain analyses to assess changes in domain-general networks, especially those dedicated to mentalizing (Amft et al.; Frith and Frith; Blakemore).

## 2. Materials and Methods

### 2.1. Participants

Participants were pairs of familiar, non-kin, agemates (± 2 years), recruited *via* web posting. They included 17 pairs of children (15/34 females) with a mean age of 11 years (range: 10-13 years) and 12 pairs of adults (16/24 females) with a mean age of 23 years (range: 20-29 years). Standardized Intellectual Quotient (IQ) was as assessed by the Nouvelle Echelle Métrique de l’Intelligence (Cognet and Bachelier) in children and by the average of the matrix reasoning and similarities subtests of the Wechsler Abbreviated Scale of Intelligence (*WAIS-IV Wechsler Adult Intelligence Scale 4th Edition*) in adults. IQs were in the normal to superior range (children, mean: 114.3, range: 76-141; adults, mean: 99.8, range: 83-115). Closeness scores, as assessed by the 7-point, Inclusion of Other in the Self scale (Aron et al.), reached scores ≥ 4 (children, mean: 5.94, range: 3-7; adults: mean: 5.54, range: 3-7), typical of close partners such as best friends (Gächter et al.; Myers and Hodges).

Nine children and one adult were discarded due to claustrophobia, sleepiness, joystick malfunction, misunderstood instructions, or excessive motion in the scanner. One of the remaining children had missing fMRI data and one of the remaining adult had missing behavioral data due to recording issues. The final samples of subjects therefore comprised 25 children and 22 adults for behavioral analyses, and 24 children and 23 adults for fMRI analyses. All participants were native French speakers, had no visual deficit, no MRI contra-indications and no history of neurological and psychiatric disorder. The study was conducted according to the guidelines of the Declaration of Helsinki, and approved by the CPP Sud Est II Ethics Committee on November 7, 2018 (ClinicalTrials.gov Identifier: NCT03453216). Informed consent was obtained from all subjects involved in the study or their parents. Each participant received a 20€-per-hour compensation for her/his time.

### 2.2. Session timeline

During the scanning session, participants first performed the numerosity comparison task (Figure 1A), and then the phonological comparison task (Figure 1B), in two successive functional runs of approximatively 12 minutes each. A pause was provided halfway through each functional run, which the participant ended at her/his convenience by pressing a button on one of the joysticks. The two functional runs were separated by an 8-minute anatomical T1 scan, and followed by a 9- minute resting state scan that is not analyzed in the present paper. Eight blocks of fixation, during which participants had to look at a fixation cross for 16,800ms, were randomly interspersed among the task blocks of each functional run.

**Figure 1:**
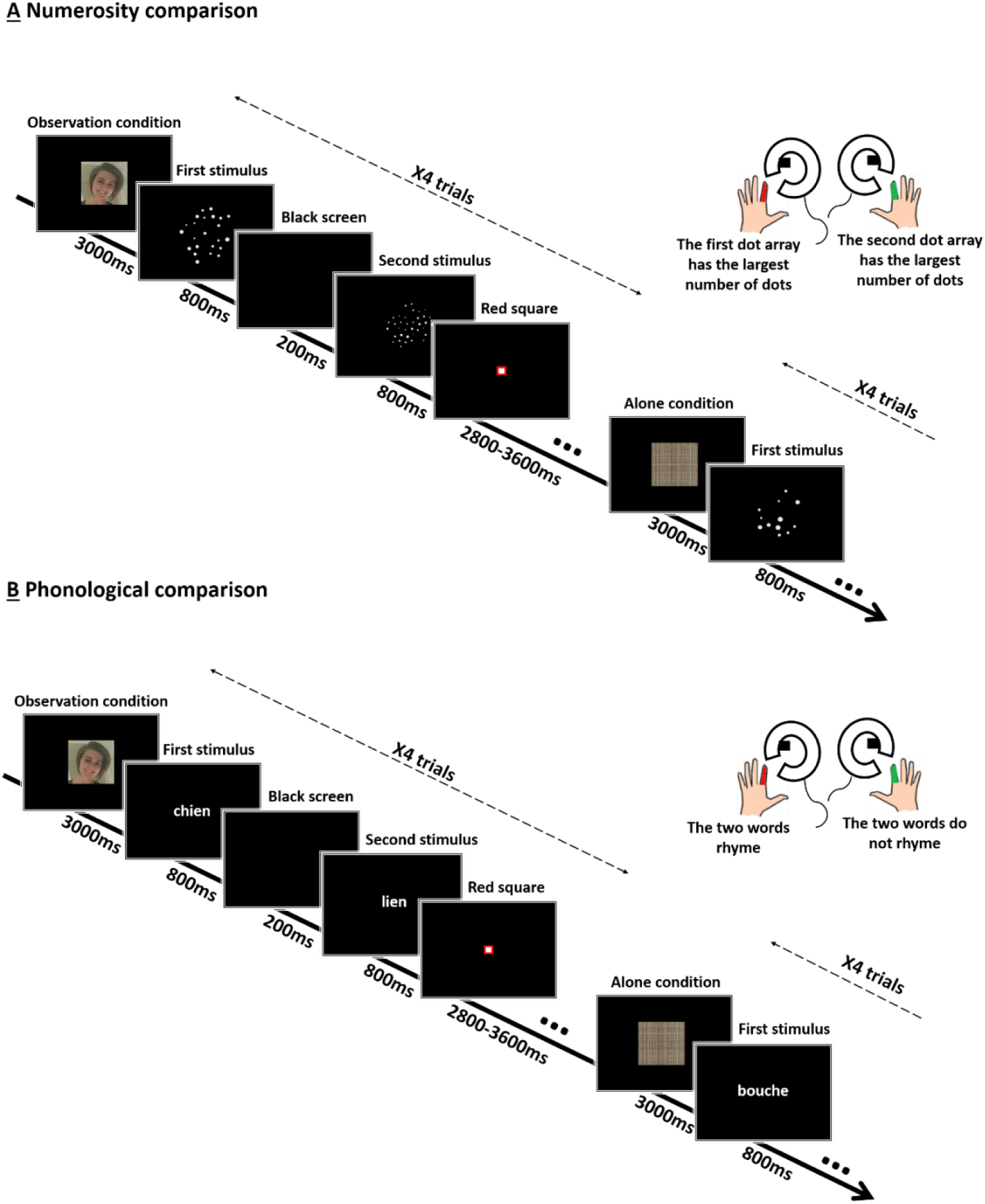
Experimental tasks. Trial time course (from top to bottom) was the same for numerosity (A) and phonological comparisons (B). At the beginning of each block of 4 trials, an original or scrambled version of a headshot of the observer indicated the condition for the block: observation or alone condition, respectively. Two stimuli were successively presented for 800ms each, separated by a 200ms-interval. Participants had to decide which of the dot arrays had the largest number of dots (A) or whether the two words rhymed or not (B), and to respond as fast and accurately as possible, using either the left or the right index finger, as soon as the second stimulus appeared and before the red square turned off.

### 2.3. Trial sequence

Both tasks were programmed using the Presentation® software (www.neurobs.comaccessed on 15 December 2021). The stimuli were projected onto a screen viewed by the participant through a mirror attached to the head coil. For each trial, two stimuli (two dot arrays for numerosity comparisons; two words for phonological comparisons) appeared one after the other for 800ms each, with a 200ms delay in between. A red square then appeared for a randomly varying duration of 2,800ms, 3,200ms or 3,600ms. Participants had to decide which array contained the largest number of dots (numerosity comparison; Figure 1A) or whether the two words rhymed or not (phonological comparison; Figure 1B). They were asked to respond as fast and accurately as possible as soon as the second stimulus appeared and before the red square turned off. Participants pressed a button of the joystick in their left hand if the first dot array had the largest number of dots or if the two words rhymed. They pressed a button of the joystick in their right hand if the second dot array had the largest number of dots, or if the two words did not rhyme.

### 2.4. Stimuli

Dot arrays were created while controlling for differences in the cumulative surface area and in the distribution of dot sizes (Gebuis and Reynvoet). An array could contain 12, 18, 24 or 36 dots. Comparison difficulty varied with the ratio of the number of dots between the two arrays. The higher the ratio, the greater the difficulty. Comparisons involved a 0.33 ratio (12 dots vs. 36 dots), a 0.5 ratio (18 dots vs. 36 dots or 12 dots vs. 24 dots), a 0.67 ratio (24 dots vs. 36 dots or 12 dots vs. 18 dots), or a 0.75 ratio (18 dots vs. 24 dots). The task was divided into 24 blocks of 4 trials each, for a total of 96 trials. There were 12 easy blocks involving small ratios (i.e., 0.33 and 0.5) and 12 difficult blocks involving large ratios (i.e., 0.67 and 0.75), pseudo-randomly ordered during the task. The first dot array contained the larger number of dots in half of the trials, while in the other half of the trials it contained the smaller number of dots. A given pair of stimuli was presented only once. The task began with 8 practice trials (4 per ratio) that were not included in the analyses.

Words contained 3 to 8 letters. Comparison difficulty varied with the congruence or incongruence of the spelling and phonology of the two successively presented words. In half of the trials, the two words had congruent orthography and phonology, i.e., they had identical spelling and sounded the same (e.g., sac-lac [sak-lak]), or they had a different spelling and sounded different (e.g., jeu-doux [ʒœ-du]). In the other half of the trials, the two words had incongruent orthography and phonology, i.e., they had a different spelling and sounded the same (e.g., dos-taux [do-to]), or they had identical spelling and sounded different (e.g., tapis-iris [tapi-iris]). The phonological comparison task was, like the numerosity comparison task, divided in 24 blocks of 4 trials each, for a total of 96 trials. There were 12 easy blocks involving congruent trials and 12 difficult blocks involving incongruent trials, pseudo-randomly ordered during the task. Each word was presented only once during the task. Words presented over two successive trials could not have the same phonology, orthography or be semantically related. The two words rhymed in half of the trials, while they did not rhyme in the other half of the trials. As for numerosity comparison, the task began with 8 practice trials (4 congruent, 4 incongruent) that were not included in the analyses.

### 2.5. Observation *versus* Alone conditions

For each pair, each subject alternatively took the actor and the observer roles. While the actor was lying inside the MR scanner, the observer was sitting in an adjacent room, facing a computer screen. The observer’s computer screen displayed filler videos (Alone condition) or the live video streams of three cameras placed inside the scanner: one filming the actor’s body, one filming the actor’s eyes, and one filming what the actor saw on her/his screen (Observation condition). The two conditions alternated every other 4-trial block, always starting with the Alone condition, up to a total of 24 blocks (96 trials) per task.

The actor was informed about the forthcoming condition at the beginning of each 4-trial block by displaying the observer’s picture for 3,000ms, either in a scrambled version (Alone condition), or in its original form (Observation condition). During the 8-minute anatomical T1 scan in between the two tasks, actor and observer could see each other via video cameras as a reminder for the actor of the observer’s actual presence in the adjacent room. During acquisition, the experimenters remained out of sight in the scanner’s monitoring room (whose window overlooking the scanner was obtruded by a curtain) and refrained from any unnecessary verbal contact with either the actor or the observer during the scanning session in order to minimize third-party presence.

### 2.6. Behavioral analyses

R (RStudio, v.1.0.136) and SYSTAT (v13) were used to analyze the subjects’ accuracy (% of correct responses) and their speed during correct responses (reaction times, RTs, calculated as the time separating the appearance of the second stimulus from the button press). Scores for each task were averaged across all 24 blocks of four trials. These averages were then analyzed using three-way ANOVAs with the between-subject factor Age (Children, Adults) and the within-subject factors Condition (Observation, Alone) and Task (Numerosity comparison, Phonological comparison). We also calculated, for each subject and each task, the performance gain produced by observation in accuracy and speed relative to the alone condition ((Observation- Alone)/Alone*100). We then used one-tailed Student’s t-tests to determine whether the group mean gain was significantly greater than 0, i.e., reflected the expected social facilitation. Peer presence effect size was estimated as earlier (Tricoche, Monfardini, et al.), using Cohen’s d (dz for dependent samples) and common language effect size (CL).

### 2.7. MRI data acquisition

MRI scans were obtained from a MAGNETOM Prisma 3.0 T scanner (Siemens Healthcare, Erlangen, Germany) at the Lyon Primage neuroimaging platform (CERMEP, Imagerie du vivant, Lyon, France). The fMRI blood oxygenation level-dependent (BOLD) signal was measured with a susceptibility weighted single-shot echo planar imaging (EPI) sequence. The following parameters were used: TR = 2,000 ms, TE = 24 ms, flip angle = 80°, matrix size = 128 x 120, field of view = 220 x 206 mm, voxel size = 1.72 x 1.72 mm, slice thickness = 3 mm (0.48 mm gap), number of slices = 32. Between the two functional runs, a high resolution T1-weighted 3D structural image was acquired for each participant (TR = 3,000 ms, TE = 2.93 ms, flip angle = 8°, matrix size = 320 x 280 mm, field of view = 280 x 320 mm, slice thickness = 0.8 mm, number of slices =160).

### 2.8. fMRI data analyses

#### 2.8.1. Preprocessing

Data analysis was performed using SPM12 (www.fil.ion.ucl.ac.uk/spm accessed on 15 September 2021). Functional images were corrected for slice acquisition delays, spatially realigned to the first image of the first run to correct for head movements, and spatially smoothed with a Gaussian filter equal to twice the voxel size (4 x 4 x 7 mm³ full width at half maximum). Functional image runs were inspected using ArtRepair (cibsr.stanford.edu/tools/human-brain-project/artrepair-software.html accessed on 15 September 2021); functional volumes with a global mean intensity greater than 3 standard deviations from the average of the run or a volume-to-volume motion greater than 2 mm were identified as outliers and substituted by the interpolation of the 2 nearest non-repaired volumes. Finally, functional images were coregistered with the segmented anatomical image and normalized into the standard Montreal Neurological Institute (MNI) space (normalized voxel size: 2 x 2 x 3.5 mm³).

#### 2.8.2. Processing

Event-related statistical analysis was conducted using the general linear model (GLM). Activation was modeled as epochs with onset time locked to the presentation of the first stimulus in each trial and with a duration of 2 seconds. Fixation periods were modeled as 16 seconds blocks. All trials (including correct, incorrect and miss trials) were sorted according to Task, Condition and trial type (e.g., ratio for numerosity comparison, congruency for phonological comparison). Fixation blocks were modeled in a separate regressor for each task. Finally, two regressors of no-interest (one per task, including instructions, breaks, and picture display of the observation/alone conditions) were added in the model. All epochs were convolved with a canonical hemodynamic response function (HRF). The time series data were high-pass filtered (1/128 Hz), and serial correlations were corrected using an autoregressive AR (1) model.

#### 2.8.3. Whole-brain analyses

Voxel-wise parameter estimates obtained for each subject were entered into random effect (RFX) analyses in order to identify regions exhibiting main effects and interactions involving the Age, Task, and/or Condition factors. Group-wise statistical maps were thresholded for significance using a voxel-wise probability threshold of p<0.001 (uncorrected) and a cluster-wise probability threshold of p<0.05 (FWE corrected for multiple comparisons).

#### 2.8.4. ROI analyses of the task-specific substrates

Task-specific ROIs were defined across groups and condition based on the main effect of task at the whole-brain level. Specifically, numerosity ROIs were defined using the contrast of numerosity comparison versus phonological comparison, while phonology ROIs were defined using the contrast of numerosity comparison versus phonological comparison. For each map, we excluded voxels for which task-related activity was not also significantly greater than activity during fixation. ROIs were defined as the intersection of 10 mm radius spheres centered on the local maximum of each cluster (using the SPM toolbox Marsbar) with the corresponding thresholded statistical map. Activity (calculated with respect to the fixation baseline) was averaged across all voxels of each ROI. These average values were then analyzed using four-way ANOVAs to assess the effects on task-specific neural activity of the between-subject factor Age (Children, Adults) and the within-subject factors Task (Numerosity comparison, Phonological comparison), Condition (Observation, Alone) and ROI. Statistical significance was set at p<0.05.

#### 2.8.5. Complementary psychophysiological interaction (PPI) analysis

Our main analyses identified a right TPJ cluster as the only area that was more activated in adults than children under peer observation (see Results section). Thus, we performed a whole-brain psycho-physiological interaction (PPI) analysis to identify brain regions whose coupling with this region was modulated as a function of Condition (Observation/Alone). The seed was defined as the entire cluster found in the Age x Condition interaction of the whole-brain analysis (coordinates of the peak: 58, -26, 10; Table 3). We estimated a GLM that included 3 regressors for each task: a physiological regressor corresponding to the entire time series of the cluster over the whole task, a psychological regressor for the Observation > Alone contrast, and the PPI regressor reflecting the interaction between the psychological and physiological regressors. The model also included two regressors of no interest (one per task).

## 3. Results

### 3.1. Behavioral data: Task and Age effects

Making phonological comparisons took longer than making numerosity comparisons (F(1,45)=177.3, p=<0.001, η_p_^2^=0.80), but accuracy was comparable in the two tasks (F(1,45)=0.2, n.s.). Children’s expectedly performed worse than adults. Their responses were less accurate (F(1,45)=5.22, p=0.03, ηp^2^ =0.10) and slower (F(1,45)=10.9, p=0.002, ηp^2^ =0.19) than adults’. Children’s developmental lag behind adults was more pronounced for phonological than for numerosity comparisons (Age x Task interaction: RTs, F(1,45)=7.73, p=0.008, ηp^2^=0.15; percent correct responses, F(1,45)=6.22, p=0.02, η ^2^=0.12).

### 3.2. Behavioral data: Condition effect

For accuracy, we found a main effect of Condition (F(1,45)=7.31, p=0.01, ηp^2^ =0.14), indicating that observation improved accuracy. This effect was qualified by a Condition x Age x Task interaction (F(1,45)=4.04, p=0.05, η ^2^=0.08), as this social facilitation was detectable (at least marginally) in all cases, i.e., children’s numerosity comparisons (Alone: 90%, Observation: 92%, gain 2%, t(24)=2.29, p=0.03), children’s phonological comparisons (Alone: 88%, Observation: 90%, gain 2%, t(24)=1.41, p=0.08), and adults’ phonological comparisons (Alone: 93%, Observation: 96%, gain 4%, t(21)=3.01, p=0.007), except adults’ numerosity comparisons (Alone: 93%, Observation: 91%). Cohen’s dz respectively estimated the three positive peer presence effects as medium- and small-sized effects of 0.52 and 0.28 for children, and as a medium-sized effect of 0.65 for adults. The corresponding CL effect sizes, which give the probability for a score randomly selected from the observed condition to be better than a score randomly selected from the unobserved condition, were: 70% and 61% for children, and 74% for adults.

For RTs, we found a Condition x Task interaction (F(1,45)=8.05, p=0.007, ηp^2^=0.15). Peer observation had no effect on phonological comparisons (Children: Alone 1324ms, Observation 1381ms; Adults: Alone 988ms, Observation 986ms), but did fasten numerosity comparisons. This held true in both children (Alone 894ms, Observation 862ms, gain 3%, t(24)=1.71, p=0.05) and adults (Alone 689ms, Observation 664ms, gain 3%, t(21)=3.24, p=0.002). Cohen’s dz estimated these peer presence effects as a small-sized effect of 0.14 for children and a medium-sized effect of 0.73 for adults. The corresponding CL effect sizes amounted to 55% for children and 77% for adults.

To summarize, behavioral data showed the expected social facilitation during observed relative to unobserved trials. Irrespective of age, the improvement under peer observation took the form of faster numerosity comparisons and more accurate phonological comparisons. Children additionally showed more accurate numerosity comparisons.

### 3.3. fMRI data: ROI analyses

The ROI analyses identified seven clusters as task-specific neural substrates (all F’s(1,135)>40, all p’s<0.001, η ^2^>0.23; Table 1, Figure 2). These clusters were consistent with earlier fMRI data obtained using the same tasks (Prado, Mutreja, Zhang, et al.; Prado, Mutreja, and Booth).

**Figure 2.**
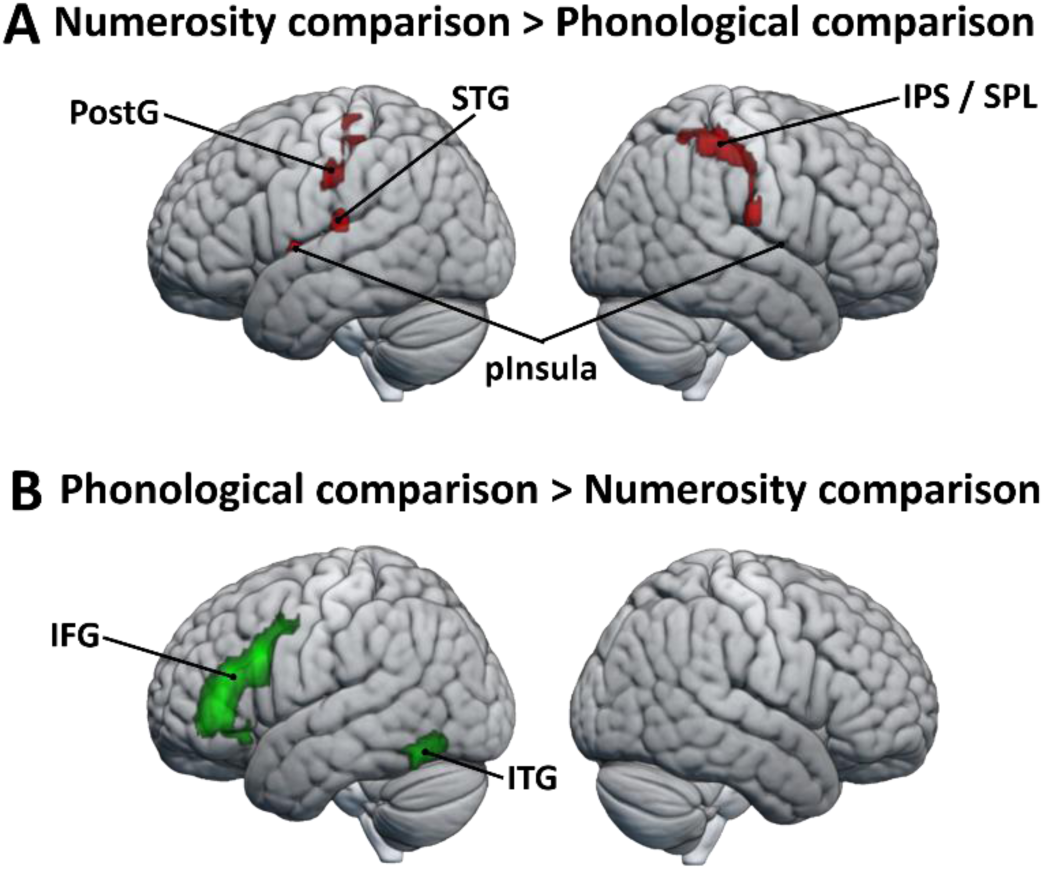
Task-specific substrates selectively activated by (A) numerosity comparisons *versus* (A) phonological comparisons.

**Table 1.** MNI (Montreal Neurological Institute) coordinates of the seven brain regions showing a main Task effect.

| Anatomical location | MNI coordinates<br>(mm) |  |  | Z score | Cluster<br>size<br>(voxels) |
| --- | --- | --- | --- | --- | --- |
|  | x | y | z |  |  |
| *Numerosity comparison > Phonological comparison* |  |  |  |  |  |
| Right Superior Parietal Lobule / Intra Parietal Sulcus | 32 | -40 | 34 | 4.94 | 616 |
| Right Posterior Insula | 46 | -4 | 13 | 4.55 | 108 |
| Left Posterior Insula | -44 | -2 | 10 | 5.45 | 207 |
| Left Superior Temporal Gyrus | -58 | -24 | 13 | 4.64 | 199 |
| Left Postcentral Gyrus | -38 | -26 | 55 | 3.93 | 94 |
| *Phonological comparison > Numerosity comparison* |  |  |  |  |  |
| Left Inferior Frontal Gyrus | -48 | 36 | 2 | 6.93 | 1244 |
| Left Inferior Temporal Gyrus | -50 | -56 | -15 | 7.06 | 317 |

There were four numerosity ROIs (see Methods): the right IPS/SPL, the right and left posterior Insula, the left STG, and the left postcentral gyrus. There were two phonological ROIs (see Methods): one in the left IFG, the other in the left ITG. A main effect of Condition was found on ROIs (F(1,1215)=4.43, p=0.03), but without any interaction with Task, Age or ROI. Yet, post-hoc tests revealed a significant Condition effect only on the left IFG (p=0.04) and a marginal effect on the left postcentral gyrus (p=0.07).

To summarize, ROI analyses provided little support to the task-specific account of peer presence effect. Considered together, ROIs showed an increase in activation under observation, but post- hoc tests revealed that this effect was significant only for one neural substrate of phonological comparisons, the left IFG. In absence of Condition x Task interaction, this increase could not be considered as reliably task-selective, i.e., as significantly greater for phonological than numerosity comparisons.

### 3.4. fMRI data: Whole-brain analyses, Condition effect

Across both groups and both tasks, no main of effect of Condition was observed in the Alone > Observation contrast. Rather, the neural correlates of peer presence effects were revealed by the whole-brain Observation > Alone contrast, which identified eleven clusters (Table 2, Figure 3A). Associated Beta values (Figure 3B) showed that the most frequent change took the form of a lesser deactivation in the observation than in the alone condition. This concerned the three nodes of the mentalizing network, the mPFC, the TPJ, and the Prec/PCC region, as well as the left MTG, right precentral gyrus PreG, and right posterior occipital gyrus (POG). Greater activation in the observation compared to the alone condition was also observed. This pattern was found in the right VS, the left IFG, the Middle Cingulate Gyrus (MCC), and a cluster involving the left frontal eye field (FEF) and extending into the precentral gyrus (PreG).

**Figure 3:**
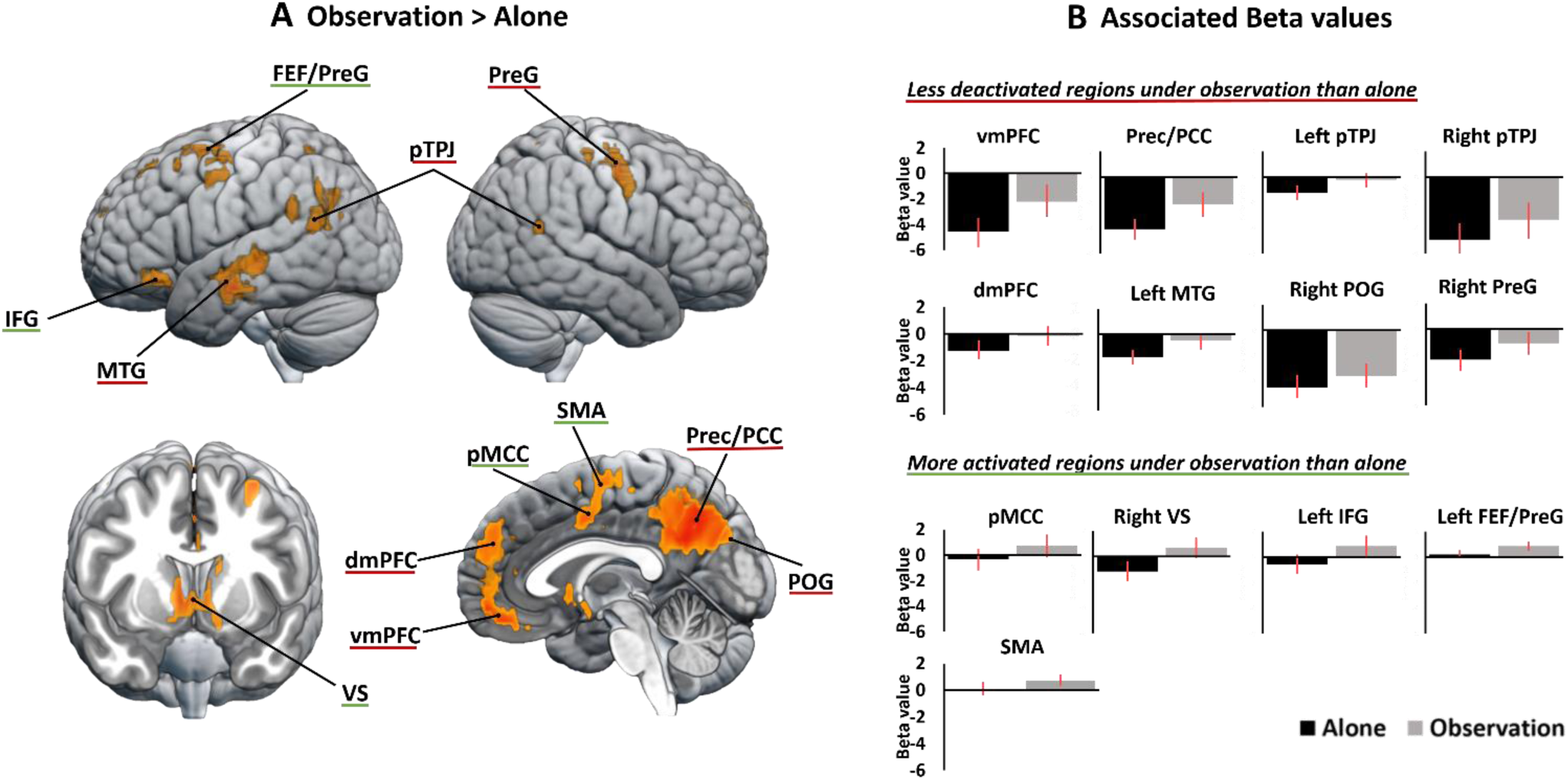
Brain regions activated in the Observation > Alone contrast across Age and Task. (A) Location of activated brain regions. (B) Associated Beta values of less deactivated regions (vmPFC, dmPFC, Prec/PCC, pTPJ, left MTG, right POG and right PreG) and more activated regions (pMCC, SMA, right VS, left IFG and left FEF/PreG) during observed relative to unobserved trials.

**Table 2.**
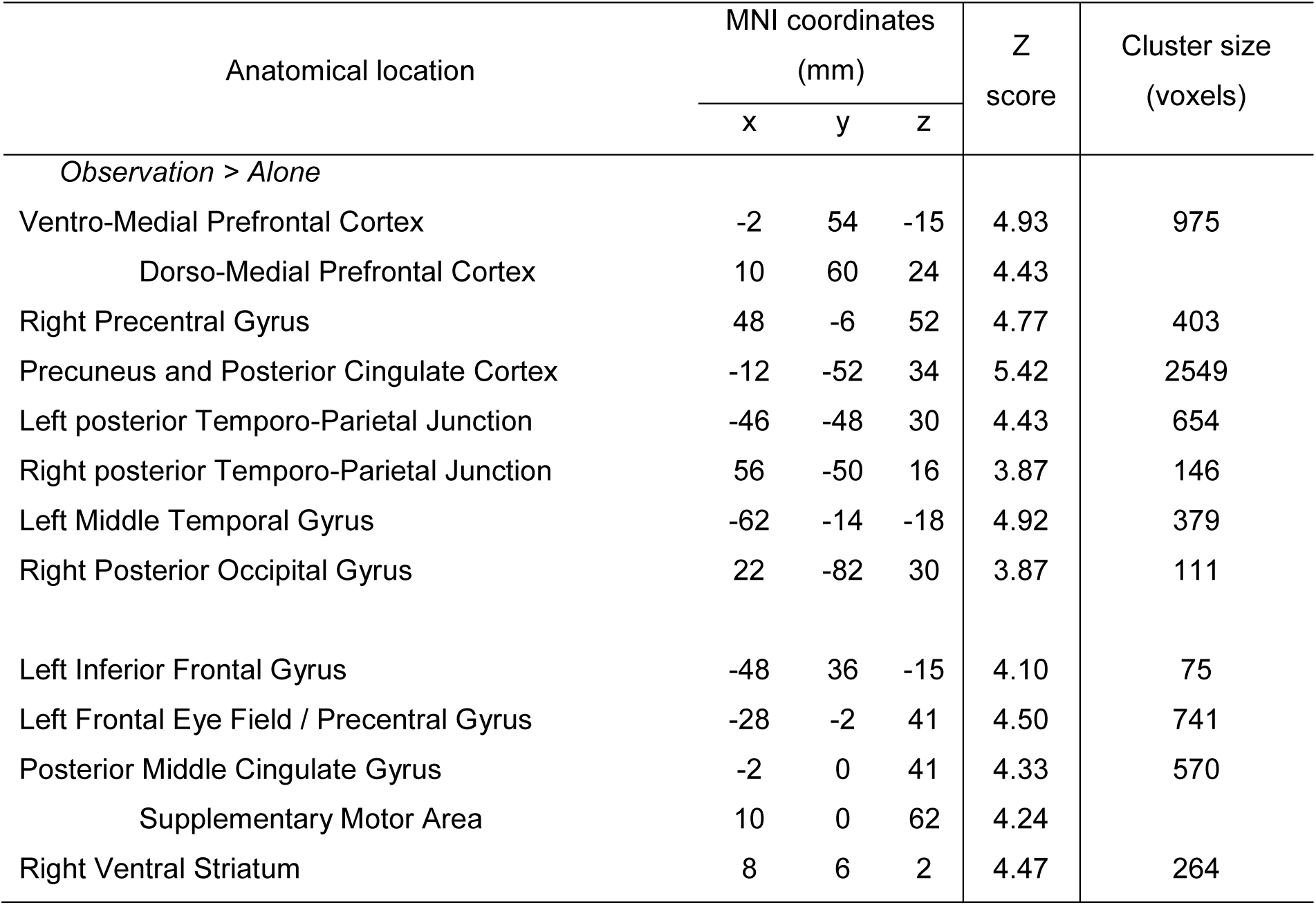
MNI coordinates of the brain regions showing a main Condition effect

### 3.5. fMRI data: Whole-brain analyses, Condition x Task interaction

Only one cluster of the large network modulated by peer observation showed a Condition x Task interaction (Figure 4). Located in the posterior part of the MCC ([2 2 41], Z=3.92), it displayed increased activation under observation for numerosity but not phonological comparisons.

**Figure 4.**
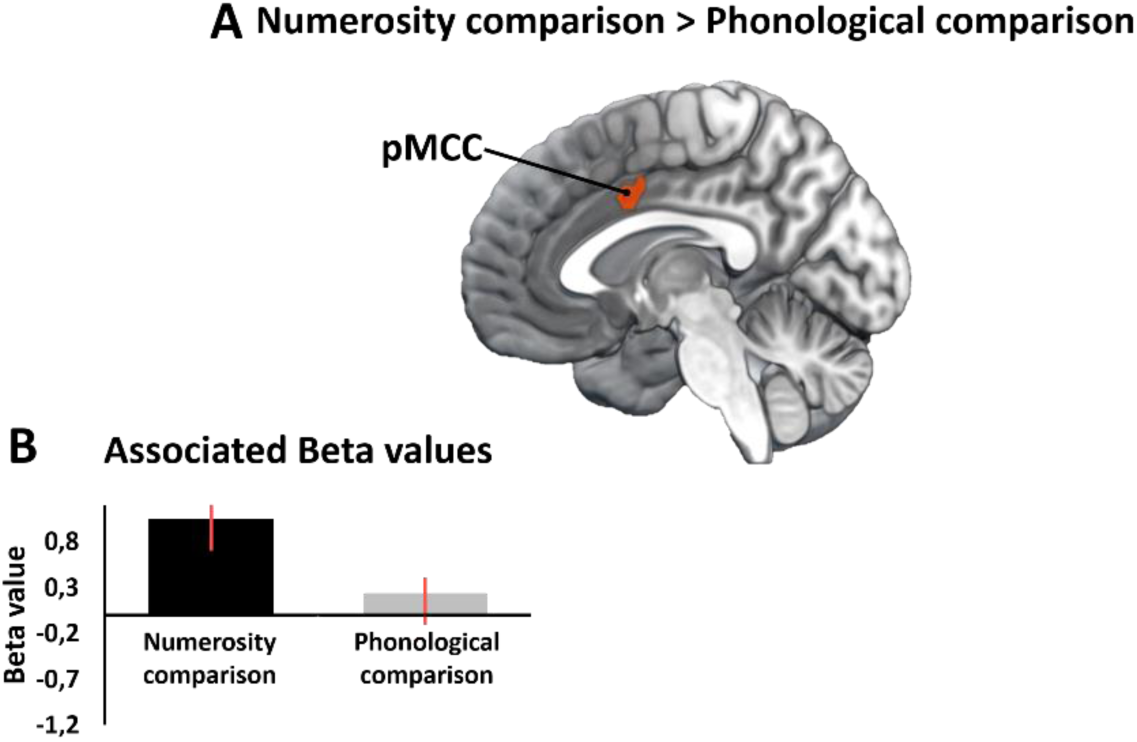
Brain activation in the Observation > Alone contrast interacting with Task. A single area (posterior MCC) showed increased activation under observation for numerosity but not phonological comparisons.

### 3.6. fMRI data: Whole-brain analyses, Condition x Age interaction

When adults were considered separately (Table 3), the brain regions identified by the Observation > Alone were the same as those described above for all subjects taken together. When children were considered separately, only one cluster survived the threshold: the Prec and nearby SPL (Figure 5B). This was also the sole cluster revealed by a conjunction analysis across adults and children for the Observation > Alone contrast (Figure 5B). However, all but one of the adults’ clusters were observed in children with a lower (p=0.005) and uncorrected statistic level (Figure 5C).

**Table 3.** MNI coordinates of the brain regions showing a main Condition effect in analyses conducted separately for Adults and Children

| Anatomical location | MNI coordinates (mm) |  |  | Z score | Cluster size (voxels) |
| --- | --- | --- | --- | --- | --- |
|  | x | y | z |  |  |
| *Observation > Alone & Adults > Children* |  |  |  |  |  |
| Right posterior Superior Temporal Sulcus (incl. anterior Temporo-Parietal Junction) | 58 | -26 | 10 | 4.79 | 126 |
| *Observation > Alone in Adults only* |  |  |  |  |  |
| Bilateral Superior Frontal cortex | 0 | 44 | 30 | 4.31 | 235 |
| (incl. medial Prefrontal Cortex) (two clusters) | 10 | 60 | 24 | 4.07 | 54 |
| Left Inferior Frontal Gyrus | -46 | 18 | 20 | 4.01 | 152 |
| Left Precentral Gyrus | -50 | -10 | 48 | 3.73 | 73 |
| Right Precentral Gyrus | 40 | -22 | 38 | 4.39 | ~~306~~7 |
| Middle Cingulate Gyrus | 0 | -6 | 30 | 4.37 | ~~445~~8 |
| Precuneus | -14 | -52 | 38 | 4.40 | ~~1439~~ |
| Right posterior Temporo-Parietal Junction | 56 | -50 | 16 | 4.19 | ~~109~~0 |
| Left anterior Temporo-Parietal Junction | -66 | -24 | -1 | 4.02 | ~~508~~1 |
| Right anterior Temporo-Parietal Junction | 60 | -26 | 13 | 4.39 | ~~242~~2 |
| Left Superior Temporal Gyrus | -50 | 0 | -4 | 4.02 | ~~122~~3 |
| Right Superior Temporal Gyrus | 46 | 4 | -15 | 3.70 | ~~567~~4 |
| Left Lingual Gyrus / Fusiform Gyrus (two clusters) | -6 | -70 | -1 | 4.35 | ~~547~~5 |
|  | -24 | -70 | -4 | 4.35 | ~~647~~6 |
| Right Ventral Striatum | 6 | 6 | -5 | 4.62 | ~~230~~7 |
|  |  |  |  |  | 478 |
| *Observation > Alone in Children only* |  |  |  |  |  |
| Precuneus | -6 | -62 | 41 | 4.29 | 260 |
| Left Superior Parietal Lobule | -14 | -56 | 62 | 4.03 | 114 |

**Figure 5:**
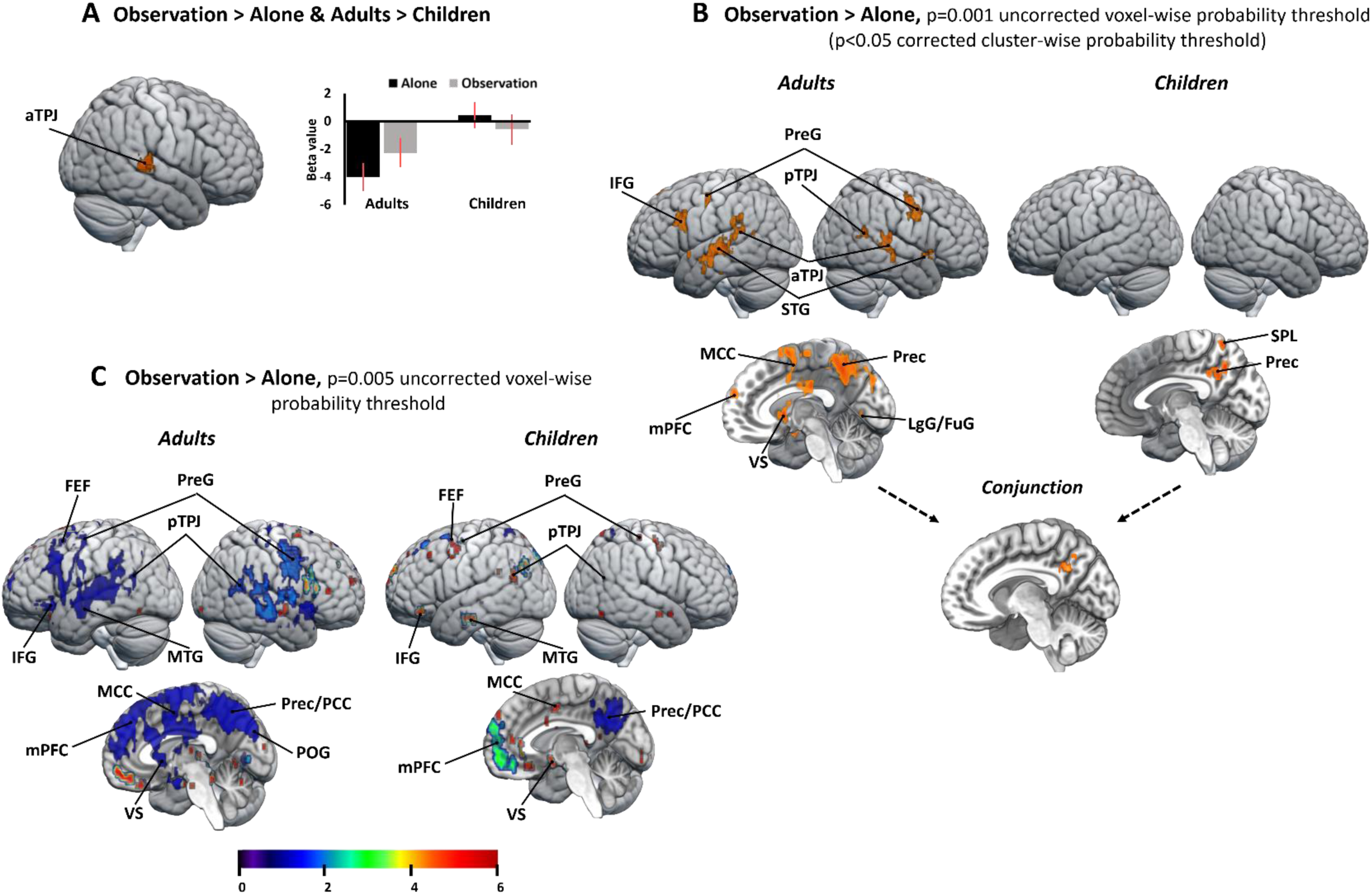
Brain activation in the Observation > Alone contrast interacting with Task (A) Brain region showing greater activity for the Observation > Alone contrast in adults than children, with associated beta values. (B) Brain regions showing greater activity for the Observation > Alone contrast in adults and children separately using a p=0.001 corrected statistic. The conjunction analysis between adults and children for the Observation > Alone contrast is shown at the bottom of the figure. (C) Brain regions showing greater activity for the Observation > Alone contrast in adults and children separately using a p=0.005 uncorrected statistic.

The one exception to the close resemblance between children and adults was a cluster located close to the right pSTS, in a region defined by Mars et al. in 2012 (Rogier B. Mars, Sallet, et al.) as the anterior part of TPJ (aTPJ, [58 -26 10], Z=4.79]). This region stood out as the only node of the large network modulated by peer observation showing a significant Condition x Age interaction. Associated Beta values showed that the right aTPJ was less deactivated in observed than unobserved trials in adults, but not in children (Figure 5A). There was no Condition x Age x Task interaction, indicating that this developmental difference held for both tasks.

### 3.7. fMRI data: PPI analysis of right aTPJ connectivity

The PPI analysis showed that peer observation decreased the connectivity of the right aTPJ cluster identified above with two visual-information-processing areas in adults, but not in children (Figure 6). These regions were the lingual gyrus (LgG, [10 -82 -8], Z=4.74) and the right [26 -70 - 12, Z=4.32] and left IOG [-34 -80 -15], Z=4.22).

**Figure 6:**
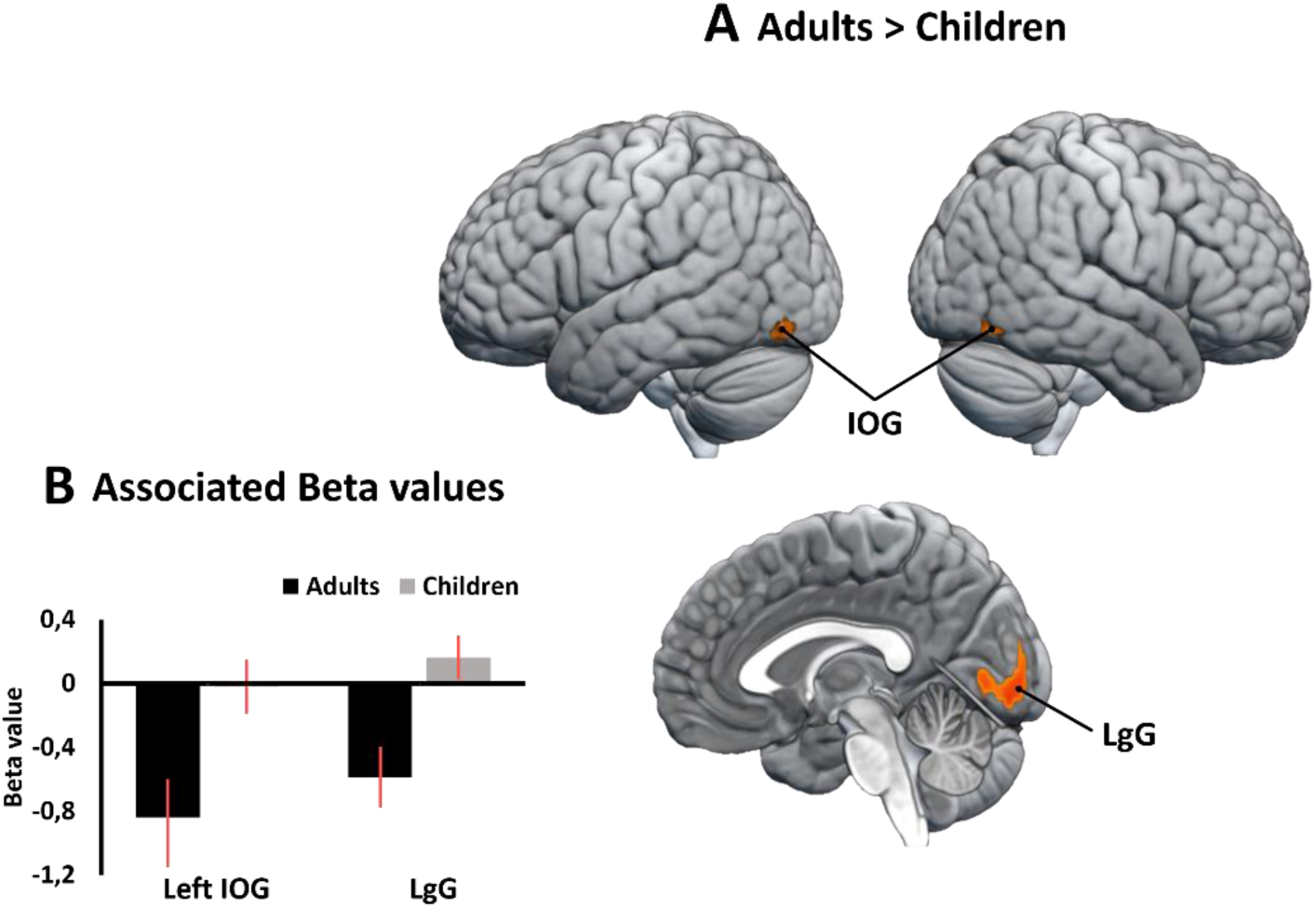
PPI analysis of the right aTPJ for the Observation > Alone contrast. (A) Brain regions showing a greater decrease of functional connectivity with the right aTPJ for the Observation than Alone condition in adults as compared to children. (B) Associated beta values (Observation > Alone contrast) measured at the peak of these regions as a function of age.

## 4. Discussion

The present study aimed to uncover the neural mechanisms underlying peer presence effects on human behavior. We investigated adults and children, alternately observed and unobserved by a familiar peer, while they compared either the number of dots in two arrays or the sounds of two written words. Behavioral findings confirmed the social facilitation of these basic skills, which are foundational to humans’ math and reading abilities. ROI analyzes revealed virtually no observation- driven change within the numerical and language brain areas forming the task-specific neural substrates of numerosity and phonological comparisons. Rather, whole-brain analyses revealed a unique neural signature of observation, similar for non-symbolic numerosities and words, and largely shared by children and adults. This task-independent signature entailed widespread changes in several brain networks known for their domain-general involvement in social cognition (especially mentalizing), attention, and reward. Children’s pattern of observation-driven neural changes largely resembled adults’, with one exception, the anterior portion of the right TPJ area. Only in adults did this area show a lesser deactivation in observed relative to unobserved trials, associated with decreased connectivity with visual-information-processing areas.

Caution should be used in generalizing the above conclusions to all peers as, here and earlier (Tricoche, Monfardini, et al.), we chose to test pairs of familiar peers. First, familiar peers are more representative of daily life as they are more frequent at school or at work than unknown ones. Second, close others capture attention (Chauhan et al.), elicit pleasure (Fareri et al.) and induce social facilitation (Herman; Sugimoto et al.; Monfardini, Reynaud, et al.) more than strangers, accordingly playing a more preeminent role in social cognition (Smith and Mackie). Notwithstanding this limitation, the present behavioral findings provide further proof of the remarkable ubiquity of social facilitation across situations and ages. First, they demonstrate that the social facilitation of numerosity and phonological comparisons reported earlier in the presence of a nearby coactor (Tricoche, Monfardini, et al.) also occurred in the less natural conditions of a live, but remote (via video) and sporadic (every other 4-trial block), observation. In our former paradigm testing numerical and phonological comparisons simultaneously, the improvement concerned speed only whereas, in the present paradigm testing numerical and phonological comparisons successively, the improvement concerned speed and accuracy. Second, the present behavioral findings show that 10 to 13-year-olds are as sensitive to social facilitation as adults, as were 8 to 10-year-olds in our previous study (Tricoche, Monfardini, et al.). Across the two studies, the magnitude of the behavioral changes produced by peer presence or peer observation in children amounted to small- to medium-sized effects (Cohen’s d: 0.14 to 0.61) not quite matching, but closely resembling those observed in adults (Cohen’s d: 0.21 to 0.73). Testing numerical and phonological comparisons in adolescents is now needed so as to unveil the full developmental trajectory of peer presence effects on these education-related skills. Being observed by a peer induces more self-conscious emotions and greater autonomic arousal in adolescents than in children and adults (Somerville et al.), and adolescence is generally viewed as the period of life with the highest susceptibility to peer influence (Albert et al.). Greater social facilitation of numerical and phonological comparisons in adolescence would indicate that peers’ influence on education-related skills follow the same inverted U-shaped trajectory as that reported for peers’ influence on reward-related behaviors (Telzer).

As detailed in the Introduction, previous studies in both human and non-human animals did describe neural changes in others’ presence in the very brain areas underlying the task at hand (Monfardini, Redoute, et al.; Dumontheil et al.; Yoshie et al.). In addition, the changes driven by peer presence in the ventral striatum, orbitofrontal cortex, and amygdala during risk-taking in adolescents can been viewed as a social modulation of the brain areas specialized in processing rewards and emotions (Chein et al.; Hoffmann et al.; Van Hoorn et al.). Yet, as most of these studies rested their conclusions on a single task, proofs of truly task-selective changes (occurring for one task but not the other) were still needed. The present study addressed this issue by using two tasks, respectively dependent on the brain numerical- and language-related areas. The regions selectively engaged by each task were consistent with earlier children’s and adults’ fMRI data collected with the same tasks (Prado, Mutreja, Zhang, et al.; Prado, Mutreja, and Booth), including, in particular, the right parietal IPS/SPS region for numerosity comparisons, and the left IFG region for phonological comparisons. ROI analyses of these clusters revealed, however, an observation-driven increase in activation in all ROIs when considered together, but with a significant effect only in the left IFG, and no observation x task interaction in any ROI. In other words, we found only weak evidence of observation-driven changes in task-specific substrates and no evidence of truly task-selective changes in support of the task-specific neural account of peer presence effects. Caution is required, however, in interpreting these negative findings. There is evidence from single-cell recordings that the primate brain harbors intertwined populations of “asocial” and “social” neurons in different brain areas. In the monkey dorsolateral prefrontal cortex, the very same task events are coded by duplicate sets of neurons: one firing when the monkey is alone, the other firing when a congener is present (Demolliens et al.). In the human dorsomedial prefrontal cortex, true and false beliefs are coded by two distinct neuronal populations, one selective for our own beliefs, the other for others’ beliefs (Jamali et al.). The local category of neurons responsible for the BOLD response being invisible to fMRI, the relative lack of evidence in support of the task-specific account of peer presence effects, including the present negative findings, might stem from limitations inherent to neuroimaging.

The present study provides, by contrast, compelling evidence in support of a domain-general neural account of peer presence effects in humans. The observation-driven changes shared across tasks and ages involved i) the right and left TPJ, a region thought to serve as a hub integrating mentalizing and higher-order attentional control (Patel et al.), together with ii) three regions known for their involvement in mentalizing: the mPFC, Prec/PCC, and left MTG (Fehlbaum et al.), and iii) three regions known for their contribution to higher-order attentional control: the left IFG, MCC, and left and right PreG/FEF (Dosenbach et al.). Observation-driven changes in the Prec/PCC cluster extended into the visual areas of the medial occipital cortex, perhaps due to our use of the peer’s picture to signal observed trials. The only other observation-driven changes shared across tasks and ages was found in the VS, a major node of the brain reward system (Haber and Knutson). In the mPFC, Prec/PCC, left MTG, and right and left TPJ, the associated beta values were negative during unobserved trials relative to the fixation baseline, and peer observation lessened this deactivation. This agrees with the fact that these four nodes of the mentalizing network overlap with the default mode network (DMN), whose trademark is to be deactivated during cognitive tasks, presumably to quiet our “default” flow of self-generated thoughts (W. Li et al.; Amft et al.; Hyatt et al.; Rogier B. Mars, Neubert, et al.). The magnitude of such task-induced deactivations tends to increase with task demands, suggesting that DMN deactivation contributes to successful task performance via an efficient reallocation of processing resources from “default” to task-relevant processes (Daselaar et al.). Lessening the magnitude of DMN deactivation necessary to achieve successful task performance could thus be one of the neural mechanisms contributing to social facilitation under peer observation. Attention studies have established that when task demands are too low, the brain becomes vulnerable to task-irrelevant stimuli; increasing task demands, in this case, improves performance by reducing or even eliminating the brain response to distractors (Lavie). Peer presence during easy tasks such as numerosity and phonological comparisons might likewise capture the resources that are left unused by the task, and dedicate them to the observer (e.g. to thoughts about her/his opinion of our performance), thereby protecting the brain from any other task-interfering distraction.

As a corollary, task-relevant stimuli might be more efficiently processed. Consistent with this hypothesis, peer observation increased activation in the dorsal frontal gyrus along the precentral sulcus, near or at the FEF, as well as in the IFG, two key nodes contributing to attentional control, usually with a predominant role of the right hemisphere (Corbetta et al.). Here, the increase concerned predominantly the left hemisphere, whose role has been less investigated. One previous study, however, showed that the left FEF and IFG form, together with the left TPJ, a pathway by which a salient contextual (task-irrelevant) cue can be translated into an attentional control signal that facilitates performance in a simple target detection task (DiQuattro and Geng). The engagement of the left FEF-IFG-TPJ pathway in the present study could reflect a similar beneficial effect of the (task-irrelevant) observer’s presence on simple responses. The increase in activation observed in a MCC cluster (extending dorsally into the SMA) might concur to improve attentional control as the MCC has been postulated as a hub implementing the higher-order attentional processes necessary for the online monitoring of responses (Procyk et al.), both our own and others’ (Apps et al.). Increasing the attentional resources dedicated to task-related information could thus be a second neural mechanism contributing to social facilitation under peer observation. Still another mechanism could be a modulation of affective valuation via the VS. Increased activity in the VS has been associated with enhanced positive valuation in others’ presence of, e.g., risk taking in adolescents (Albert et al.), or monetary gain in adults (Fareri et al.). In the present study, no feedback was provided to participants about their accuracy and no reward (praise or money) was given for correct responses. The VS increase in activation under peer observation might therefore reflects an enhancement of the reward intrinsic to live social interactions (Pfeiffer et al.).

Overall, the present study provides evidence that peer observation (remote, and episodic, but live), in absence of any reward (save the presence of a friend) and any explicit mentalizing demands, nevertheless triggers widespread neural changes combining attenuated deactivation of mentalizing/DMN nodes with enhanced activation of attentional control and reward-related regions. These findings are consistent with earlier descriptions of the remarkable power of live social interaction on our brain. Specifically, extensive neural changes reminiscent of the present ones, i.e., also including mentalizing, attention, and reward regions, have been reported previously in both adults and children who are (or believe they are) engaged in a real-time interaction with a social (unknown) partner, instead of either watching a recorded version of the same social interaction, or having the same real-time interaction with a computer (Redcay et al.; Rice Warnell et al.; Rice et al.). The engagement of the mentalizing network during peer observation and live social interaction might result, in both cases, from spontaneous mentalizing, that is, from wondering about the observer’s thoughts, or a readiness to do so at any moment (Merchant et al.; Hamilton and Lind). Yet, the present observation-driven changes extend beyond the mentalizing network, and live social interaction was previously shown to elicit more extensive activation, both within and outside the mentalizing network, than tasks explicitly requiring to infer another’s mental state (Alkire et al.; Rice and Redcay). So, spontaneous conscious mentalizing alone seems insufficient to explain the widespread changes produced by peer observation or live social interaction. Evidence is growing that, starting from infancy, others’ representations of the world are automatically coded by our brain, even when irrelevant to our goal, and influence our responses without our awareness (Kampis and Southgate; Steinmetz and Pfattheicher; Smith and Mackie). The resources taken up by such automatic coding inevitably alters the balance between task-dedicated and default mode networks, as well as the distribution of attentional resources to vs. away from the task. This could explain why the mere presence of a social partner can simultaneously affect multiple networks including, in humans, social cognition, attention, and reward, simply by forcing the brain to change the way it harnesses its limited resources.

Regarding development, the present study provides new insights into children’s peer presence effects, which have been much less investigated than adults’. A conjunction analysis identified the Prec/PCC as the region presenting the most robust observation-driven change shared by children and adults. This region is a core mentalizing node (Schurz et al.). It stands out, in addition, as a unique hub distinguishing between task and rest states in both the developing and adult human brain (R. Li et al.). Its engagement in the present 10 to 13-year-olds is in agreement with previous findings demonstrating that precuneal mentalizing and DMN networks are already present and functional by the time children reach school age (R. Li et al.). The conjunction analysis did not identify the other observation-driven changes identified in adults at the p<0.001 corrected level (TPJ, mPFC, Prec/PCC, MTG, IFG, MCC, and PreG/FEF) as shared with children. None of these clusters showed, however, a reliable Age x Condition interaction. In fact, all of them were present in children at a more lenient p<0.005 uncorrected statistical threshold, likely resulting from the development noisiness that typically makes children’s behavior more variable and neural networks less functional specialized than adults’ (Richardson et al.). Thus, although development likely sharpens them thereafter (see below), brain changes mediating peer presence effects are already present and functional by 10-13 years of age. This neural resemblance of children and adults parallels the comparable behavioral magnitude of peer presence effects that we observed here and earlier (Tricoche, Monfardini, et al.) across childhood and adulthood.

An abundant developmental literature has established that children’s mentalizing capacities, despite their early presence in life, do undergo continuous refinement over development, evolving from relatively simple insights into others’ perceptions and goals in infants and toddlers to a sophisticated understanding of others’ sarcasm and irony in early adolescence (Richardson et al.). Given its dependence upon the brain mentalizing networks, response to others’ presence likely undergoes similar refinement over development. This hypothesis is corroborated by the results of the present Age x Condition interaction analysis, which did identify one reliable age-dependent difference in observation-driven neural changes. It concerned the right TPJ, a region associated with attention and social cognition (Carter and Huettel), two intertwined functions as early attentional capacities predict later social cognition abilities during development (Mundy and Newell). In adults, there is evidence that the right TPJ harbors two functional entities: a posterior region exclusively dedicated to social cognition, especially mentalizing, and an anterior region, bridging attention and social cognition through its role in shifting attention between both stimuli and mental states (Krall et al.; Rogier B. Mars, Sallet, et al.). In the present study, observation-driven changes encompassed both TPJ regions, but the cluster identified by the interaction analysis as a specifically adult change seem to more closely correspond to the anterior portion of rTPJ. This raises the possibility that, because of its sophisticated integrative functions, the anterior portion of rTPJ stands out as having not yet reached its mature role in the control of peer presence effects by late childhood.

## 5. Conclusions

The present study tested the hypothesis that peer presence effects rely on a neural combination of task-selective changes and domain-general modulations using a developmental approach comparing children to adults. The results did not reveal any reliable task-selective changes, but the possibility remains that such changes occur at levels invisible to fMRI. They did, by contrast, provide compelling evidence for widespread task-independent changes in domain-general brain networks that are already in place in late childhood. Putting together phylogenetic and ontogenetic perspectives is the challenge awaiting future studies in order to explain the neural implementation of all social presence effects, from the rudimentary ones shared by infants and animals to the most sophisticated ones that are the privilege of healthy human adults.

## Acknowledgments

We thank the CERMEP Primage MRI engineers Franck Lamberton, and Danielle Ibarrola for invaluable help in data collection. We thank the Centre Hospitalier le Vinatier for the promotion of this study. We thank Jérôme Redouté for his support throughout the study and his comments on the manuscript, and Fadila Hadj-Bouziane for helpful discussions. We extend our thanks to all the volunteers who took part in the study, with special thanks to the children.

## Data Availability

Data supporting the results will be available on the Open Science Framework website at https://osf.io/nspkx/.

## References

1. Albert, Dustin, et al. “The Teenage Brain: Peer Influences on Adolescent Decision Making.” Current Directions in Psychological Science, vol. 22, no. 2, 2013, pp. 114–20, doi:10.1177/0963721412471347.

2. Alkire, Diana, et al. “Social Interaction Recruits Mentalizing and Reward Systems in Middle Childhood.” Human Brain Mapping, vol. 39, no. 10, 2018, pp. 3928–42, doi:10.1002/hbm.24221.

3. Amft, Maren, et al. “Definition and Characterization of an Extended Social-Affective Default Network.” Brain Structure and Function, vol. 220, no. 2, 2015, pp. 1031–49, doi:10.1007/s00429-013-0698-0.

4. Apps, Matthew A. J., et al. “The Role of the Midcingulate Cortex in Monitoring Others’ Decisions.” Frontiers in Neuroscience, vol. 7, no. 7 DEC, 2013, pp. 1–7, doi:10.3389/fnins.2013.00251.

5. Aron, A., et al. “Inclusion of Other in the Self Scale and the Structure of Interpersonal Closeness.” Journal of Personality and Social Psychology, vol. 63, no. 4, 1992, doi:https://doi.org/10.1037/0022-3514.63.4.596.

6. Belletier, Clément, et al. “Social-Facilitation-and-Impairment Effects: From Motivation to Cognition and the Social Brain.” Current Directions in Psychological Science, vol. 28, no. 3, June 2019, pp. 260–65, doi:10.1177/0963721419829699.

7. Beyer, Frederike, et al. “Losing Control in Social Situations: How the Presence of Others Affects Neural Processes Related to Sense of Agency.” Eneuro, vol. 5, no. 1, 2018, p. ENEURO.0336-17.2018, doi:10.1523/ENEURO.0336-17.2018.

8. Blakemore, Sarah Jayne. “Development of the Social Brain in Adolescence.” Self-Regulation in Adolescence, 2015, pp. 193–211, doi:10.1017/CBO9781139565790.010.

9. Bond, and Linda J. Titus. “Social Facilitation: A Meta-Analysis of 241 Studies.” Psychological Bulletin, vol. 94, no. 2, Sept. 1983, pp. 265–92, doi:https://doi.org/10.1037/0033-2909.94.2.265.

10. Carter, R. Mc Kell, and Scott A. Huettel. “A Nexus Model of the Temporal-Parietal Junction.” Trends in Cognitive Sciences, vol. 17, no. 7, Elsevier Ltd, 2013, pp. 328–36, doi:10.1016/j.tics.2013.05.007.

11. Chabaud, Marie-Ange, et al. “Social Facilitation of Long-Lasting Memory Retrieval in Drosophila.” Current Biology : CB, vol. 19, no. 19, Elsevier Ltd, Oct. 2009, pp. 1654–59, doi:10.1016/j.cub.2009.08.017.

12. Chauhan, Vassiki, et al. “Social Saliency of the Cue Slows Attention Shifts.” Frontiers in Psychology, vol. 8, no. MAY, 2017, pp. 1–11, doi:10.3389/fpsyg.2017.00738.

13. Chein, Jason, et al. “Peers Increase Adolescent Risk Taking by Enhancing Activity in the Brain’s Reward Circuitry.” Developmental Science, vol. 14, no. 2, 2011, pp. 1–10, doi:10.1111/j.1467-7687.2010.01035.x.Peers.

14. Chib, Vikram S., et al. “Neural Substrates of Social Facilitation Effects on Incentive-Based Performance.” Social Cognitive and Affective Neuroscience, vol. 13, no. 4, 2018, pp. 391–403, doi:10.1093/scan/nsy024.

15. Cognet, Georges, and Delphine Bachelier. NEMI-2 Nouvelle Échelle Métrique de l’intelligence – Deuxième Version. Edited by « Les outils du psychologue », Dunod, 2017, https://www.cairn.info/clinique-de-l-examen-psychologique-de-l-enfant--9782100763658-page-331.htm.

16. Corbetta, Maurizio, et al. “The Reorienting System of the Human Brain: From Environment to Theory of Mind.” Neuron, vol. 58, no. 3, 2008, pp. 306–24, doi:10.1016/j.neuron.2008.04.017.

17. Daselaar, S. M., et al. “When Less Means More: Deactivations during Encoding That Predict Subsequent Memory.” NeuroImage, vol. 23, no. 3, 2004, pp. 921–27, doi:10.1016/j.neuroimage.2004.07.031.

18. Dehaene, Stanislas. “Précis of ‘ The Number Sense .’” Mind and Language, vol. Wiley Onli, 2001, pp. 1–24, doi:10.1111/j.1468-0017.1986.tb00086.x.

19. Demolliens, Marie, et al. “Social and Asocial Neurons in the Prefrontal Cortex : A New Look on Social Facilitation and the Social Brain .” Social Cognitive and Affective Neuroscience, vol. 12, no. 8, 2017, pp. 1241–48, doi:10.1093/scan/nsx053.

20. DiQuattro, Nicholas E., and Joy J. Geng. “Contextual Knowledge Configures Attentional Control Networks.” Journal of Neuroscience, vol. 31, no. 49, 2011, pp. 18026–35, doi:10.1523/JNEUROSCI.4040-11.2011.

21. Dosenbach, Nico U. F., et al. “A Core System for the Implementation of Task Sets.” Neuron, vol. 50, no. 5, 2006, pp. 799–812, doi:10.1016/j.neuron.2006.04.031.

22. Dumontheil, Iroise, et al. “Audience Effects on the Neural Correlates of Relational Reasoning in Adolescence.” Neuropsychologia, vol. 87, Elsevier, 2016, pp. 85–95, doi:10.1016/j.neuropsychologia.2016.05.001.

23. Ehri, Linnea C., et al. “Phonemic Awareness Instruction Helps Children Learn to Read: Evidence From the National Reading Panel’s Meta-Analysis.” Reading Research Quarterly, vol. 36, no. 3, 2001, pp. 250–87, doi:10.1598/rrq.36.3.2.

24. Fareri, Dominic S., et al. “Social Network Modulation of Reward-Related Signals.” The Journal of Neuroscience : The Official Journal of the Society for Neuroscience, vol. 32, no. 26, June 2012, pp. 9045–52, doi:10.1523/JNEUROSCI.0610-12.2012.

25. Fehlbaum, Lynn V, et al. “Early and Late Neural Correlates of Mentalizing: ALE Meta-Analyses in Adults, Children and Adolescents.” Social Cognitive and Affective Neuroscience, no. July 2020, 2021, pp. 1–16, doi:10.1093/scan/nsab105.

26. Ferrari, Pier F., et al. “Mirror Neurons through the Lens of Epigenetics.” Trends in Cognitive Sciences, vol. 17, no. 9, 2013, pp. 450–57, doi:10.1016/j.tics.2013.07.003.

27. Frith, Chris D., and Uta Frith. “The Neural Basis of Mentalizing.” Neuron, vol. 50, no. 4, 2006, pp. 531–34, doi:10.1016/j.neuron.2006.05.001.

28. Gächter, Simon, et al. “Measuring the Closeness of Relationships: A Comprehensive Evaluation of the ‘inclusion of the Other in the Self’ Scale.” PLoS ONE, vol. 10, no. 6, 2015, pp. 1–19, doi:10.1371/journal.pone.0129478.

29. Gebuis, Titia, and Bert Reynvoet. “Generating Nonsymbolic Number Stimuli.” Behavior Research Methods, vol. 43, no. 4, 2011, pp. 981–86, doi:10.3758/s13428-011-0097-5.

30. Guerin, Bernard. Social Facilitation. Cambridge University Press, 2010.

31. Haber, Suzanne N., and Brian Knutson. “The Reward Circuit: Linking Primate Anatomy and Human Imaging.” Neuropsychopharmacology : Official Publication of the American College of Neuropsychopharmacology, vol. 35, no. 1, Nature Publishing Group, Jan. 2010, pp. 4–26, doi:10.1038/npp.2009.129.

32. Hamilton, Antonia F. de C., and Frida Lind. “Audience Effects: What Can They Tell Us about Social Neuroscience, Theory of Mind and Autism?” Culture and Brain, vol. 4, no. 2, Springer Berlin Heidelberg, 2016, pp. 159–77, doi:10.1007/s40167-016-0044-5.

33. Hartley, Catherine A., and Leah H. Somerville. “The Neuroscience of Adolescent Decision- Making.” Current Opinion in Behavioral Sciences, vol. 5, 2015, pp. 108–15, doi:10.1016/j.cobeha.2015.09.004.

34. Hazem, Nesrine, et al. “I Know You Can See Me: Social Attention Influences Bodily Self- Awareness.” Biological Psychology, vol. 124, Elsevier B.V., Mar. 2017, pp. 21–29, doi:10.1016/j.biopsycho.2017.01.007.

35. Herman, C. Peter. “The Social Facilitation of Eating. A Review.” Appetite, vol. 86, Elsevier Ltd, 2015, pp. 61–73, doi:10.1016/j.appet.2014.09.016.

36. Hessler, N. A., and A. J. Doupe. “Social Context Modulates Singing-Related Neural Activity in the Songbird Forebrain.” Nature Neuroscience, vol. 2, no. 3, 1999, pp. 209–11, doi:10.1038/6306.

37. Higgs, Suzanne, and Jason Thomas. “Social Influences on Eating.” Current Opinion in Behavioral Sciences, vol. 9, Elsevier Ltd, 1 June 2016, pp. 1–6, doi:10.1016/j.cobeha.2015.10.005.

38. Hoffmann, Ferdinand, et al. “Risk-Taking, Peer-Influence and Child Maltreatment: A Neurocognitive Investigation.” Social Cognitive and Affective Neuroscience, vol. 13, no. 1, Oxford University Press, Jan. 2018, pp. 124–34, doi:10.1093/scan/nsx124.

39. Huguet, Pascal, et al. “Cognitive Control under Social Influence in Baboons.” Journal of Experiment Psychology: General, vol. 143, no. 6, 2014, pp. 2067–73, http://psycnet.apa.org/journals/xge/143/6/2067/.

40. Hyatt, Christopher J., et al. “Specific Default Mode Subnetworks Support Mentalizing as Revealed through Opposing Network Recruitment by Social and Semantic FMRI Tasks.” Human Brain Mapping, vol. 36, no. 8, 2015, pp. 3047–63, doi:10.1002/hbm.22827.

41. Hyde, Daniel C., et al. “Brief Non-Symbolic, Approximate Number Practice Enhances Subsequent Exact Symbolic Arithmetic in Children.” Cognition, vol. 131, no. 1, Elsevier B.V., 2014, pp. 92–107, doi:10.1016/j.cognition.2013.12.007.

42. Jamali, Mohsen, et al. “Single-Neuronal Predictions of Others’ Beliefs in Humans.” Nature, vol. 591, no. 7851, Nature Research, Mar. 2021, pp. 610–14, doi:10.1038/s41586-021-03184-0.

43. Kampis, Dora, and Victoria Southgate. “Altercentric Cognition: How Others Influence Our Cognitive Processing.” Trends in Cognitive Sciences, vol. 24, no. 11, 2020, pp. 945–59, doi:10.1016/j.tics.2020.09.003.

44. Kovács, Ágnes Melinda, et al. “Can Infants Adopt Underspecified Contents into Attributed Beliefs? Representational Prerequisites of Theory of Mind.” Cognition, vol. 213, 2021, doi:10.1016/j.cognition.2021.104640.

45. Krall, S. C., et al. “The Role of the Right Temporoparietal Junction in Attention and Social Interaction as Revealed by ALE Meta-Analysis.” Brain Structure and Function, vol. 220, no. 2, 2015, pp. 587–604, doi:10.1007/s00429-014-0803-z.

46. Lavie, N. “Attention, Distraction, and Cognitive Control Under Load.” Current Directions in Psychological Science, vol. 19, no. 3, June 2010, pp. 143–48, doi:10.1177/0963721410370295.

47. Li, Rosa, et al. “Developmental Maturation of the Precuneus as a Functional Core of the Default Mode Network.” Journal of Cognitive Neuroscience, vol. 31, no. 10, 2019, pp. 1506–19, doi:https://doi.org/10.1162/jocn_a_01426G.

48. Li, Wanqing, et al. “The Default Mode Network and Social Understanding of Others: What Do Brain Connectivity Studies Tell Us.” Frontiers in Human Neuroscience, vol. 8, no. 1 FEB, 2014, pp. 1–15, doi:10.3389/fnhum.2014.00074.

49. Liu, Na, and Ruifeng Yu. “Influence of Social Presence on Eye Movements in Visual Search Tasks.” Ergonomics, vol. 60, no. 12, Taylor & Francis, 2017, pp. 1667–81, doi:10.1080/00140139.2017.1342870.

50. Mars, Rogier B., Jérôme Sallet, et al. “Connectivity-Based Subdivisions of the Human Right ‘Temporoparietal Junction Area’: Evidence for Different Areas Participating in Different Cortical Networks.” Cerebral Cortex, vol. 22, no. 8, 2012, pp. 1894–903, doi:10.1093/cercor/bhr268.

51. Mars, Rogier B, et al. “On the Relationship between the ‘ Default Mode Network ’ and the ‘ Social Brain .’” Frontiers in Human Neuroscience, vol. 6, no. June, 2012, pp. 1–9, doi:10.3389/fnhum.2012.00189.

52. Mars, Rogier B., Franz Xaver Neubert, et al. “On the Relationship between the ‘Default Mode Network’ and the ‘Social Brain.’” Frontiers in Human Neuroscience, vol. 6, no. JUNE 2012, 2012, pp. 1–9, doi:10.3389/fnhum.2012.00189.

53. Menardy, F., et al. “The Presence of an Audience Modulates Responses to Familiar Call Stimuli in the Male Zebra Finch Forebrain.” European Journal of Neuroscience, no. April, 2014, pp. 1–13, doi:10.1111/ejn.12696.

54. Merchant, Junaid S., et al. “Neural Similarity between Mentalizing and Live Social Interaction.” Preprint, 2021.

55. Miyazaki, Yuki. “Influence of Being Videotaped on the Prevalence Effect during Visual Search.” Frontiers in Psychology, vol. 6, no. May, 2015, pp. 1–7, doi:10.3389/fpsyg.2015.00583.

56. Monfardini, Elisabetta, J. Redoute, et al. “Others’ Sheer Presence Boosts Brain Activity in the Attention (But Not the Motivation) Network.” Cerebral Cortex, 2015, pp. 1–13, doi:10.1093/cercor/bhv067.

57. Monfardini, Elisabetta, Amélie J. Reynaud, et al. “Social Modulation of Cognition: Lessons from Rhesus Macaques Relevant to Education.” Neuroscience and Biobehavioral Reviews, vol. 82, 2017, pp. 45–57, doi:10.1016/j.neubiorev.2016.12.002.

58. Müller-Pinzler, L., et al. “Neural Pathways of Embarrassment and Their Modulation by Social Anxiety.” NeuroImage, vol. 119, Elsevier B.V., 2015, pp. 252–61, doi:10.1016/j.neuroimage.2015.06.036.

59. Mundy, Peter, and Lisa Newell. Attention, Social Joint Cognition Attention,. no. 5, 2013, pp. 269–74.

60. Muria, Aurélie, et al. “Social Facilitation of Long-Lasting Memory Is Mediated by CO2 in Drosophila.” Current Biology, vol. 31, no. 10, 2021, pp. 2065–2074.e5, doi:10.1016/j.cub.2021.02.044.

61. Myers, Michael W., and Sara D. Hodges. “The Structure of Self-Other Overlap and Its Relationship to Perspective Taking.” Personal Relationships, vol. 19, no. 4, 2012, pp. 663–79, doi:10.1111/j.1475-6811.2011.01382.x.

62. Patel, Gaurav H., et al. “The Evolution of the Temporoparietal Junction and Posterior Superior Temporal Sulcus.” Cortex, vol. 118, 2019, pp. 38–50, doi:10.1016/j.cortex.2019.01.026.

63. Pearcey, Sharon M., and John M. D. E. Castro. “Food Intake and Meal Patterns of One Year Old Infants.” Appetite, vol. 29, 1997, pp. 201–12.

64. Pfeiffer, Ulrich J., et al. “Why We Interact: On the Functional Role of the Striatum in the Subjective Experience of Social Interaction.” NeuroImage, vol. 101, 2014, pp. 124–37, doi:10.1016/j.neuroimage.2014.06.061.

65. Phillips, Beth M., et al. “Successful Phonological Awareness Instruction With Preschool Children: Lessons From the Classroom.” Topics in Early Childhood Special Education, vol. 28, no. 1, 2008, pp. 3–17, doi:10.1177/0271121407313813.

66. Prado, Jérôme, Rachna Mutreja, and James R. Booth. “Developmental Dissociation in the Neural Responses to Simple Multiplication and Subtraction Problems.” Developmental Science, vol. 17, no. 4, 2014, pp. 537–52, doi:10.1111/desc.12140.

67. Prado, Jérôme, Rachna Mutreja, Hongchuan Zhang, et al. “Distinct Representations of Subtraction and Multiplication in the Neural Systems for Numerosity and Language.” Human Brain Mapping, vol. 32, no. 11, 2011, pp. 1932–47, doi:10.1002/hbm.21159.

68. Preckel, Katrin, et al. “On the Interaction of Social Affect and Cognition: Empathy, Compassion and Theory of Mind.” Current Opinion in Behavioral Sciences, vol. 19, Elsevier Ltd, Feb. 2018, pp. 1–6, doi:10.1016/j.cobeha.2017.07.010.

69. Procyk, Emmanuel, et al. “Midcingulate Motor Map and Feedback Detection: Converging Data from Humans and Monkeys.” Cerebral Cortex, vol. 26, no. 2, 2016, pp. 467–76, doi:10.1093/cercor/bhu213.

70. Qi, Sharon, and Rollanda O’Connor. “Comparison of Phonological Training Procedures in Kindergarten Classrooms.” Journal of Educational Research, vol. 93, no. 4, 2000, pp. 226–33, doi:10.1080/00220670009598711.

71. Redcay, Elizabeth, et al. “Live Face-to-Face Interaction during FMRI: A New Tool for Social Cognitive Neuroscience.” NeuroImage, 2010, doi:10.1016/j.neuroimage.2010.01.052.

72. Reynaud, Amélie J., et al. “Social Facilitation of Cognition in Rhesus Monkeys: Audience Vs. Coaction.” Frontiers in Behavioral Neuroscience, vol. 9, no. December, 2015, pp. 1–5, doi:10.3389/fnbeh.2015.00328.

73. Rice, Katherine, et al. “Perceived Live Interaction Modulates the Developing Social Brain.” Social Cognitive and Affective Neuroscience, vol. 11, no. 9, 2016, pp. 1354–62, doi:10.1093/scan/nsw060.

74. Rice, Katherine, and Elizabeth Redcay. “Interaction Matters: A Perceived Social Partner Alters the Neural Processing of Human Speech.” NeuroImage, vol. 129, 2016, doi:10.1016/j.neuroimage.2015.11.041.

75. Rice Warnell, Katherine, et al. “Let’s Chat: Developmental Neural Bases of Social Motivation during Real-Time Peer Interaction.” Developmental Science, vol. 21, no. 3, 2018, pp. 1–14, doi:10.1111/desc.12581.

76. Richardson, Hilary, et al. “Development of the Social Brain from Age Three to Twelve Years.” Nature Communications, vol. 9, no. 1, Springer US, 2018, pp. 1–12, doi:10.1038/s41467-018-03399-2.

77. Riters, Lauren V., et al. “Vocal Production in Different Social Contexts Relates to Variation in Immediate Early Gene Immunoreactivity within and Outside of the Song Control System.” Behavioural Brain Research, vol. 155, no. 2, 2004, pp. 307–18, doi:10.1016/j.bbr.2004.05.002.

78. Ruddock, Helen K., et al. “A Systematic Review and Meta-Analysis of the Social Facilitation of Eating.” American Journal of Clinical Nutrition, vol. 110, no. 4, 2019, pp. 842–61, doi:10.1093/ajcn/nqz155.

79. Schurz, Matthias, et al. “Fractionating Theory of Mind: A Meta-Analysis of Functional Brain Imaging Studies.” Neuroscience and Biobehavioral Reviews, vol. 42, Elsevier Ltd, 2014, pp. 9–34, doi:10.1016/j.neubiorev.2014.01.009.

80. Smith, Ashley R., et al. “Peers Influence Adolescent Reward Processing, but Not Response Inhibition.” Cognitive, Affective and Behavioral Neuroscience, vol. 18, no. 2, Springer New York LLC, Apr. 2018, pp. 284–95, doi:10.3758/s13415-018-0569-5.

81. Smith, Eliot R., and Diane M. Mackie. “Influence from Representations of Others’ Responses: Social Priming Meets Social Influence.” Current Opinion in Psychology, vol. 12, Elsevier Ltd, 2016, pp. 22–25, doi:10.1016/j.copsyc.2016.04.012.

82. Somerville, Leah H. “Sensitivity to Social Evaluation.” Current Directions in Psychological Science, vol. 22, no. 2, 2013, pp. 121–27, doi:10.1177/0963721413476512.

83. Somerville, Leah H. “The Medial Prefrontal Cortex and the Emergence of Self-Conscious Emotion in Adolescence.” Psychol Sci, vol. 24, no. 8, 2013, pp. 1554–62, doi:10.1177/0956797613475633.

84. Starr, Ariel, et al. “Number Sense in Infancy Predicts Mathematical Abilities in Childhood.” Proceedings of the National Academy of Sciences of the United States of America, vol. 110, no. 45, 2013, pp. 18116–20, doi:10.1073/pnas.1302751110.

85. Steinberg, Laurence. “Risk Taking in Adolescence: New Perspectives from Brain and Behavioral Science.” Current Directions in Psychological Science, vol. 16, no. 2, 2007, pp. 55–59, doi:10.1111/j.1467-8721.2007.00475.x.

86. Steinmetz, Janina, and Stefan Pfattheicher. “Beyond Social Facilitation: A Review of the Far- Reaching Effects of Social Attention.” Social Cognition, vol. 35, no. 5, 2017, pp. 585–99, doi:10.1521/soco.2017.35.5.585.

87. Sugimoto, Hikaru, et al. “Competing against a Familiar Friend: Interactive Mechanism of the Temporo-Parietal Junction with the Reward-Related Regions during Episodic Encoding.” *NeuroImage*, Elsevier B.V., 2016, doi:10.1016/j.neuroimage.2016.02.020.

88. Telzer, Eva H. “Dopaminergic Reward Sensitivity Can Promote Adolescent Health: A New Perspective on the Mechanism of Ventral Striatum Activation.” Developmental Cognitive Neuroscience, vol. 17, Elsevier Ltd, Feb. 2016, pp. 57–67, doi:10.1016/j.dcn.2015.10.010.

89. Telzer, Eva H., Nicholas T. Ichien, et al. “Mothers Know Best: Redirecting Adolescent Reward Sensitivity toward Safe Behavior during Risk Taking.” Social Cognitive and Affective Neuroscience, vol. 10, no. 10, Oxford University Press, Feb. 2015, pp. 1383–91, doi:10.1093/scan/nsv026.

90. Telzer, Eva H., Christina R. Rogers, et al. “Neural Correlates of Social Influence on Risk Taking and Substance Use in Adolescents.” Current Addiction Reports, vol. 4, no. 3, Springer Nature, Sept. 2017, pp. 333–41, doi:10.1007/s40429-017-0164-9.

91. Tricoche, Leslie, Elisabetta Monfardini, et al. “Peer Presence Effect on Numerosity and Phonological Comparisons i in 4th Graders: When Working With a Schoolmate Makes Children More Adult-Like.” Biology, vol. 10, no. 902, 2021, doi:10.3390/biology10090902.

92. Tricoche, Leslie, Johan Ferrand-Verdejo, et al. “Peer Presence Effects on Eye Movements and Attentional Performance.” Frontiers in Behavioral Neuroscience, vol. 13, no. January, 2020, pp. 1–13, doi:10.3389/fnbeh.2019.00280.

93. van Duijvenvoorde, Anna C. K., et al. “What Motivates Adolescents? Neural Responses to Rewards and Their Influence on Adolescents’ Risk Taking, Learning, and Cognitive Control.” Neuroscience and Biobehavioral Reviews, vol. 70, 2016, pp. 135–47, doi:10.1016/j.neubiorev.2016.06.037.

94. van Hoorn, Jorien, et al. “Differential Effects of Parent and Peer Presence on Neural Correlates of Risk Taking in Adolescence.” Social Cognitive and Affective Neuroscience, vol. 13, no. 9, Oxford University Press, 2018, pp. 945–55, doi:10.1093/scan/nsy071.

95. Van Hoorn, Jorien, et al. “Neural Correlates of Prosocial Peer Influence on Public Goods Game Donations during Adolescence.” Social Cognitive and Affective Neuroscience, 2016, p. nsw013, doi:10.1093/scan/nsw013.

96. van Meurs, Edda, et al. “Moving in the Presence of Others – a Systematic Review and Meta- Analysis on Social Facilitation.” PsyArXiv Preprints, vol. 3, no. 2, 2021, p. 6.

97. van Veluw, Susanne J., and Steven A. Chance. “Differentiating between Self and Others: An ALE Meta-Analysis of FMRI Studies of Self-Recognition and Theory of Mind.” Brain Imaging and Behavior, vol. 8, no. 1, 2014, pp. 24–38, doi:10.1007/s11682-013-9266-8.

98. Vignal, Clémentine, Nicolas Mathevon, et al. “Audience Drives Male Songbird Response to Partner’s Voice.” Nature, vol. 430, no. 6998, 2004, pp. 448–51, doi:10.1038/nature02645.

99. Vignal, Clémentine, Julie Andru, et al. “Social Context Modulates Behavioural and Brain Immediate Early Gene Responses to Sound in Male Songbird.” The European Journal of Neuroscience, vol. 22, no. 4, Aug. 2005, pp. 949–55, doi:10.1111/j.1460-9568.2005.04254.x.

100. WAIS-IV Wechsler Adult Intelligence Scale 4th Edition.

101. Woods, Sarah, et al. “Is Someone Watching Me? - Consideration of Social Facilitation Effects in Human-Robot Interaction Experiments.” *Proceedings of IEEE International Symposium on Computational Intelligence in Robotics and Automation*, CIRA, 2005, pp. 53–60, doi:10.1109/cira.2005.1554254.

102. Woolley, Sarah C., et al. “Emergence of Context-Dependent Variability across a Basal Ganglia Network.” Neuron, vol. 82, no. 1, Elsevier Inc., Apr. 2014, pp. 208–23, doi:10.1016/j.neuron.2014.01.039.

103. Woolley, Sarah C. “Social Context Differentially Modulates Activity of Two Interneuron Populations in an Avian Basal Ganglia Nucleus.” Journal of Neurophysiology, vol. 116, no. 6, 2016, pp. 2831–40, doi:10.1152/jn.00622.2016.

104. Wykowska, Agnieszka, et al. “Beliefs about the Minds of Others Influence How We Process Sensory Information.” PLoS ONE, vol. 9, no. 4, 2014, p. 94339, doi:10.1371/journal.pone.0094339.

105. Xin, Qiuhong, et al. “Selective Contribution of the Telencephalic Arcopallium to the Social Facilitation of Foraging Efforts in the Domestic Chick.” European Journal of Neuroscience, vol. 45, no. 3, 2017, pp. 365–80, doi:10.1111/ejn.13475.

106. Yoshie, Michiko, et al. “Why I Tense up When You Watch Me: Inferior Parietal Cortex Mediates an Audience’s Influence on Motor Performance.” Scientific Reports, vol. 6, Nature Publishing Group, 2016, p. 19305, doi:10.1038/srep19305.

107. Zajonc, RB. “Social Facilitation.” Science, vol. 149, 1965, pp. 269–74, doi:10.1126/science.149.3681.269.

